# DNA methylation mediates genetic liability to non-syndromic cleft lip/palate

**DOI:** 10.1101/256842

**Authors:** Laurence J Howe, Tom G Richardson, Ryan Arathimos, Lucas Alvizi, Maria-Rita Passos-Bueno, Philip Stanier, Ellen Nohr, Kerstin U Ludwig, Elisabeth Mangold, Michael Knapp, Evie Stergiakouli, Beate St Pourcain, George Davey Smith, Jonathan Sandy, Caroline L Relton, Sarah J Lewis, Gibran Hemani, Gemma C Sharp

## Abstract

**Background:** Non-syndromic cleft lip/palate (nsCL/P) is a complex trait with genetic and environmental risk factors. Around 40 distinct genetic risk loci have been identified for nsCL/P, but many reside in non-protein-coding regions with an unclear function. We hypothesised that one possibility is that the genetic risk variants influence susceptibility to nsCL/P through gene regulation pathways, such as those involving DNA methylation.

**Methods:** Using nsCL/P Genome-wide association study summary data and methylation data from four studies, we used Mendelian randomization and joint likelihood mapping to identify putative loci where genetic liability to nsCL/P may be mediated by variation in DNA methylation in blood.

**Results:** There was evidence at three independent loci, *VAX1* (*10q25.3*), *LOC146880* (17q23.3) and *NTN1* (17p13.1), that liability to nsCL/P and variation in DNA methylation might be driven by the same genetic variant. Follow up analyses using DNA methylation data, derived from lip and palate tissue, and gene expression catalogues provided further insight into possible biological mechanisms.

**Conclusions:** Genetic variation may increase liability to nsCL/P by influencing DNA methylation and gene expression at *VAX1, LOC146880* and *NTN1*.

## Introduction

Orofacial clefts are a heterogenous group of birth disorders ^1^. In epidemiology and genetics research, orofacial clefts can be divided into the subtypes cleft palate only (CPO) and cleft lip with or without cleft palate (CL/P), with strong evidence for distinct aetiologies ^1^. There is also accumulating evidence suggesting that the CL/P subtypes cleft lip only (CLO) and cleft lip with cleft palate (CLP) may also differ aetiologically ^2 3^. Mendelian syndromes can feature CL/P and CPO but around 70% of CL/P cases are non-syndromic (nsCL/P), with a complex aetiology likely to involve both genetic and environmental risk factors ^4 5^.

Genome-wide association studies (GWAS) have identified around 40 distinct genetic risk variants for nsCL/P in European and Asian populations ^2 6-13^ but many variants reside in non-protein coding regions and so their functional relevance remains unclear. One possibility is that genetic risk variants may be affecting nsCL/P susceptibility through gene regulation pathways. Indeed, a non-coding interval at 8q24, a major nsCL/P risk locus, has previously been shown to regulate gene expression in the developing murine face ^14^. There is increasing evidence that epigenetic mechanisms, such as DNA methylation, play a role in development of orofacial clefts ^15-18^, potentially via changes to gene expression.

In this study, we applied a recently devised analysis framework ^19 20^ to explore whether genetic influences on liability to nsCL/P are mediated by DNA methylation assayed in whole blood (Figure 1). Genetic variants that are associated with DNA methylation (methylation quantitative trait loci; mQTLs) have been previously identified in the Avon Longitudinal Study of Parents and Children (ALSPAC)^21^. We use these mQTLs to perform Mendelian randomization (MR), an epidemiological tool typically used to explore causal relationships between modifiable risk factors and health or disease outcomes ^22^. In this instance, genetic variants robustly associated with DNA methylation were tested for association with nsCL/P. We considered four possible models to explain an association between an mQTL and nsCL/P: 1) DNA methylation mediates genetic influences on liability to nsCL/P; 2) the direction of effect is reversed, i.e. genetic liability to nsCL/P causes variation in DNA methylation; 3) DNA methylation and liability to nsCL/P are influenced by separate genetic variants that are in linkage disequilibrium (LD) with each other; 4) DNA methylation and liability to nsCL/P are influenced by the same genetic variant, but via independent pathways, i.e. the association is due to horizontal pleiotropy (Figure 1)^19 20^.

**Figure 1.**
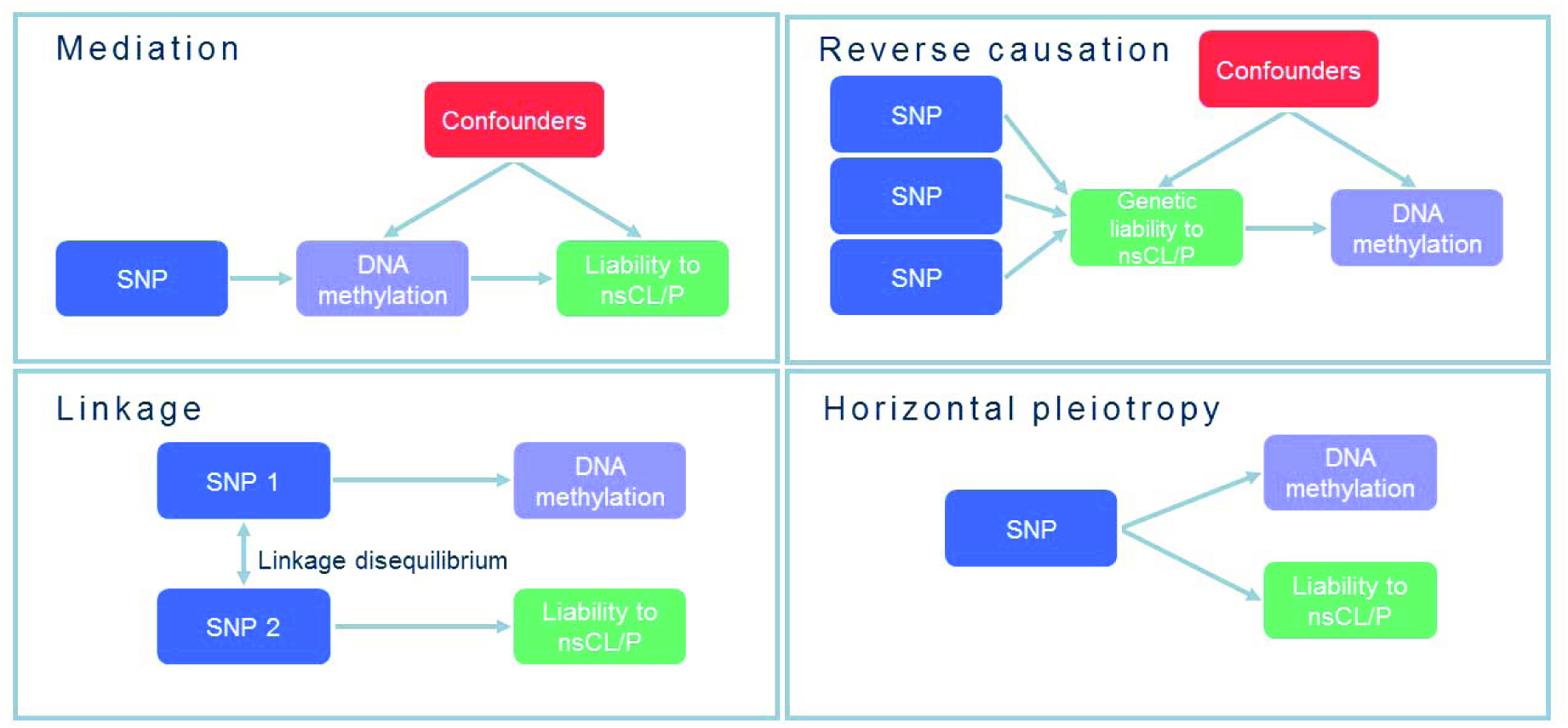
Possible explanations for an association between a methylation quantitative trait loci (mQTL) and nsCL/P. In this paper, we attempt to identify loci where genetic influences on nsCL/P are mediated by DNA methylation, i.e. the top left-hand box.

We systematically applied additional analyses, including bidirectional MR and co-localization, to estimate the most likely model as far as possible although we were unable to use recently derived MR methods ^23 24^ to distinguish between mediation and vertical or horizontal pleiotropy because most CpGs are instrumented by a single genetic variant. Where there was evidence that genetic influences on liability to nsCL/P may be mediated by DNA methylation, we explored associations with gene expression. We also compared our findings from the general population to results from an epigenome-wide association study (EWAS) of whole blood samples from nsCL/P cases and unaffected controls. Given accumulating evidence that different subtypes of OFCs have distinct aetiologies, we also explored whether identified CpGs are differentially methylated in blood samples from children with different OFC subtypes. Finally, since the majority of our analyses used DNA methylation derived in blood, which might not be representative of the developing orofacial tissues, as described previously ^15^, we explored correlations between DNA methylation in blood and lip/palate tissue in the same individuals.

## >Methods

### Data sources

#### nsCL/P genetic risk variants

We identified single nucleotide polymorphisms (SNPs) associated with nsCL/P by conducting a meta-analysis of summary statistics from two nsCL/P GWAS. Summary statistics for the first GWAS came from a case-control study of 399 cases and 1318 controls of Central European descent ^7^. For the second GWAS, we generated summary statistics by conducting a GWAS using individual level data from 638 parent-offspring trios and 178 parent-offspring duos of European descent from the International Consortium to Identify Genes and Interactions Controlling Oral Clefts (ICC). These data were available to download from dbGaP (Study Accession phs000094.v1.p1) ^25^. Full GWAS methods are described in the **Supplementary Material**, but briefly, we performed a transmission disequilibrium test (TDT) ^26^ on the pedigree data. We then performed a fixed-effects inverse-variance-weighted meta-analysis of the summary statistics from both GWAS using METAL, on the total sample of 1215 cases and 2772 controls ^27^. The results compared well with those previously published using a very similar dataset but slightly different quality control and analysis methods ^6^. We used LiftOver (genome.sph.umich.edu/wiki/LiftOver) to convert the genome positions in the nsCL/P summary statistics to the most recent genome build 37. Finally, we used PLINK ^28^ and ALSPAC as a reference panel clump the results according to LD (r^2^<0.001), within a 250 kb region around each index variant, and generate a set of independent SNPs for the pipeline.

#### Methylation genetic risk variants (mQTLs)

##### ALSPAC

To identify mQTLs (SNPs associated with DNA methylation), we used data from the Avon Longitudinal Study of Parents and Children (ALSPAC) ^29 30^. In addition to collecting detailed questionnaire and clinic data for the whole cohort, the study has generated genome-wide DNA methylation and genotype data for subsets of the cohort ^31^ (methods described in the **Supplementary Material**). These data have previously been used to generate a database of mQTLs (http://www.mqtldb.org/)^21^. The database contains summary statistics for all mQTLs with a P-value <1x10^-7^ for the association between SNP and CpG. For the purposes of this study, we focused on the mQTLs identified in cord blood samples collected at birth (the closest available time point to the orofacial developmental period). For part of our study (the reverse two sample MR), we required specific CpG-SNP associations that were unavailable from mQTLdb.org. Therefore, for required CpGs, we replicated the methods in the original study: we excluded individuals with missing genotype or covariate data, leaving 787 children. We then rank-normalised the methylation data to remove outliers and controlled for covariates, potential batch effects and the influence of cell heterogeneity by regressing data points on sex, the first 10 ancestry principal components, bisulfite-converted DNA batch and blood cell proportions estimated using the Houseman method ^32 33^. We then calculated residuals, which were used as the outcome variable in a linear regression model in PLINK^28^ to calculate the relevant CpG-SNP associations.

Finally, we excluded any mQTLs acting in trans (i.e. any SNP associated with a CpG site more than 1M base pairs away) and excluded any CpGs that have been flagged as potentially problematic (for example, cross-hybridising probes) according to a previous publication ^34^.

##### GOYA

We attempted to replicate mQTLs from ALSPAC using genotype and cord blood DNA methylation data from the Genetics of Overweight Young Adults (GOYA) cohort, which is a subset of the Danish National Birth Cohort (DNBC) ^35^. Genotype and cord blood DNA methylation data were available for 1000 children. We replicated the methods described above for ALSPAC by first excluding individuals with missing genotype or covariate data, leaving 889 children and also removing SNPs with missingness (>5%) using PLINK. As in ALSPAC, we rank-normalised the methylation data to remove outliers and adjusted for covariates, potential batch effects and the influence of cell heterogeneity by regressing data points on sex, the first 10 ancestry principal components, DNA batch and blood cell proportions estimated using the Houseman method ^32 33^. Residuals were then used as the outcome variable in a linear regression model in PLINK to calculate the relevant CpG-SNP associations.

#### Expression quantitative trait loci (eQTLs) as genetic risk variants *GTEx*

To identify eQTLs (SNPs associated with gene expression), we used Genotype-Tissue Expression (GTEx, www.gtexportal.org), which is a database of eQTLs generated using genotype and RNA sequencing gene expression data for 43 distinct tissue types from 175 individuals ^36 37^.

#### NESDA NTR Conditional eQTL Catalog

To explore the consistency of our findings, we also identified eQTLs using a second database: the NESDA NTR Conditional eQTL Catalog (https://eqtl.onderzoek.io/index.php?page=info). For this database, eQTLs were identified using genotype and gene expression microarray data from blood samples from 4896 individuals across two Dutch biobanks. Conditional eQTL analysis was applied to distinguish between dependent and independent eQTLs ^38^.

#### DNA methylation in children with orofacial clefts *Brazilian cohort*

To assess whether methylation at nsCL/P-associated CpGs (identified through MR) differs between nsCL/P cases and controls, we performed a look-up of results from a recently-published EWAS^15^. This EWAS compared blood DNA methylation profiles in 67 non-familial, nsCL/P cases and 59 age-and sex-matched controls from a Brazilian population. The average age at sampling was 5.29 years for cases and 6.45 years for controls. DNA methylation was measured using the Illumina Infinium HumanMethylation450 BeadChip platform.

#### The Cleft Collective

To explore whether methylation at nsCL/P-associated CpGs differs by cleft subtype, we compared mean methylation values in blood and matched lip/palate tissue samples from 150 children from the UK enrolled in the Cleft Collective birth cohort study. Methylation data were generated for a separate study, as previously described^2^. Briefly, a sample of 150 believed-to-be-non-syndromic children was randomly selected and stratified by cleft subtype: 50 with cleft lip only (CLO), 50 with cleft palate only (CPO), and 50 with cleft lip and palate (CLP). These children have been classified as non-syndromic because they have not been diagnosed as having any other anomaly, however, since the children are still very young, we cannot be completely sure of their non-syndromic status. Blood and either lip or palate tissue samples were available for each of the 150 children in this study. The orofacial tissue type was dependent on the OFC subtype; therefore, lip samples were available for children with CLO and palate samples for children with CPO. Of the 50 children with CLP, 43 contributed a lip sample and seven contributed a palate sample. Genome-wide DNA methylation was measured using the Illumina Infinium HumanMethylation450 BeadChip platform and functional normalisation was performed on the blood and tissue samples together. Of the original 300 samples, three blood and two lip samples failed quality control. Surrogate variables were generated using the sva package in R to capture variation in the methylation data associated with technical batch and cellular heterogeneity ^39^.

### Analysis pipeline

#### Testing for mediation: Mendelian randomization of the effect of methylation on liability to nsCL/P

nsCL/P meta-GWAS summary statistics for 543,150 SNPs were LD-pruned (r^2^<0.001) to 17,090 independent SNPs. These independent SNPs were then merged with 127,215 mQTLs from the ALSPAC mQTL database. After removing potentially problematic CpGs and CpGs acting in trans (which may increase the likelihood of horizontal pleiotropy), there were 7,091 independent CpG-SNP pairings for 6,425 distinct CpGs. We then used the MR-base R package^40^ to perform two-sample MR on all CpGs, using mQTLs as the exposure variables and nsCL/P as the outcome. In initial analysis, CpGs with one mQTL were tested using the Wald test and CpGs with more than one mQTL were tested using the Inverse Variance Weighted (IVW) method. To account for possible residual LD between mQTLs, CpGs with more than one mQTL, were retested adjusting for LD between the SNPs using a likelihood-based method ^41^. Pair-wise SNP LD was computed using the Caucasian European (CEU) and British (GBR) populations in LDlink^42^. As a sensitivity analysis, we attempted to replicate the SNP-CpG associations with a Bonferroni-corrected MR p-value <0.05 in GOYA.

#### Testing for reverse causation: Mendelian randomization of the effect of genetic liability to nsCL/P on methylation

To assess the possibility of reverse causation, we used MR-base to conduct the reverse two sample MR. Six genome-wide significant nsCL/P SNPs in Europeans ^6^ were used as the exposure and mQTLs from ALSPAC were used as the outcome. The IVW method was used as the primary analysis.

#### Testing for linkage: joint-likelihood mapping to assess co-localisation

We used the Joint Likelihood Mapping (JLIM) package in R (jlim.R)^43^ to test if liability to nsCL/P and methylation are driven by the same causal effect in each region of interest, i.e. the mQTL and nsCL/P risk variant are co-localised rather than simply being in LD. To distinguish between separate causal variants, we set the limit of genetic resolution in terms of r^2^ to 0.8. 1000 genomes CEU data was used as the reference dataset for LD. Most CpGs were associated with only one independent mQTL, so we were not able to distinguish mediation/vertical pleiotropy (top left-hand panel of **Figure 1**) from horizontal pleiotropy (bottom right-hand panel of **Figure 1**).

#### Comparison to gene expression

The previous steps identified CpGs that potentially mediate the effect of genetic variation on susceptibility to nsCL/P. Further evidence for a functional effect would be provided if mQTLs also affected gene expression. Therefore, we looked up relevant SNPs in two eQTL databases (GTEx ^37^ and NESDA NTR Conditional eQTL Catalog ^38^) and noted the estimated effect size and P-values for eQTLs in various tissues.

#### Comparison to EWAS results

At identified CpGs, we looked-up the mean methylation values in nsCL/P cases and controls from the Brazilian EWAS study. We compared the direction of estimated effect and P-values obtained using the observational EWAS and MR approaches.

#### Tissue and cleft-subtype-specific variation

At identified CpGs, we used data from the Cleft Collective to explore 1) whether methylation in blood was correlated with methylation at the site of the cleft (lip/palate), and 2) whether mean methylation varied according to cleft subtype (CLO, CPO or CLP). One-way ANOVA was used to compare the mean methylation of subtypes, adjusting for sex and surrogate variables designed to capture technical batch and cell composition effects.

## Results

### Testing for mediation: Mendelian randomization of the effect of methylation on liability to nsCL/P

To identify CpGs where methylation may mediate genetic liability to nsCL/P, we used two sample MR with mQTLs from the ALSPAC data as the exposure and liability to nsCL/P as the outcome (from the nsCL/P GWAS meta-analysis summary statistics). We found evidence for an effect of methylation on liability to nsCL/P at 26 CpGs after Bonferroni correction for 6,425 tests (Bonferroni-corrected P-value <0.05, corresponding to an uncorrected P-value <7.8x10^-6^). Of these 26 CpGs, 20 were instrumented by single mQTLs and six were instrumented by two mQTLs each. When the six CpGs with two mQTLs each were re-tested taking into account LD between the SNPs, only one (cg02598441 at *LOC146880*) survived correction for multiple testing. These 21 mQTLs were therefore taken forward to the reverse-causation step.

As a sensitivity analysis, we investigated all 21 of the ALSPAC mQTLs in data from the GOYA cohort. 17 of the 21 CpG-SNP pairings passed quality control and were present in the GOYA data, of which 16 replicated in the same direction with P<0.05 (**Supplementary Table 1**).

### Testing for reverse causation: Mendelian randomization of the effect of genetic liability to nsCL/P on methylation

Next, we tested if the association between the mQTLs and liability to nsCL/P arose because genetic liability to nsCL/P influences variation in methylation, by using two sample MR with genetic liability to nsCL/P as the exposure and methylation as the outcome. We found no evidence that genetic liability to nsCL/P influences variation in methylation at the 21 CpGs (**Table 1**). However, it should be noted that this step is very likely to be limited by statistical power

**Table 1.** Results of the forward (methylation → nsCL/P) and reverse (nsCL/P → methylation) Mendelian randomisation and the co-localisation analyses in ALSPAC.

| **SNP (allele 1/allele 2; annotated gene)** | **CpG (annotated gene)** | **Forward MR (effect size [standard error]; P-value)** | **Reverse MR (effect size [standard error]; P-value)** | **Co-localisation (JLIM statistic; P-value by permutation)** |
| --- | --- | --- | --- | --- |
| rs12057415 (T/C; n/a) | cg09549015 (*F3*) | -1.1 [0.2]; $1.1 \times 10^{-6}$ | 0.04 [0.07]; 0.59 | -8.5; 1 |
| rs12057415 (T/C; n/a) | cg26112574 (n/a) | 0.7 [0.1]; $1.1 \times 10^{-6}$ | 0.00 [0.05]; 1.00 | -16.7; 1 |
| rs861020 (A/G; *IRF6*) | cg12766975 (*IRF6*) | 1.1 [0.2]; $1.1 \times 10^{-6}$ | 0.00 [0.04]; 0.99 | -11.8; 1 |
| rs861020 (A/G; *IRF6*) | cg09163369 (*C1orf107*) | -0.5 [0.1]; $1.1 \times 10^{-6}$ | -0.04 [0.08]; 0.59 | -36.1; 1 |
| rs861020 (A/G; *IRF6*) | cg23166289 (*C1orf107*) | -0.7 [0.2]; $1.1 \times 10^{-6}$ | -0.01 [0.06]; 0.92 | -36.8; 1 |
| rs861020 (A/G; *IRF6*) | cg05527609 (*C1orf107*) | 0.9 [0.2]; $1.1 \times 10^{-6}$ | -0.06 [0.06]; 0.31 | -2.1; 0.69 |
| rs4422741 (C/T; n/a) | ch.8.2579072R (n/a) | 0.7 [0.1]; $2.1 \times 10^{-10}$ | 0.11 [0.06]; 0.08 | -2.4; 0.91 |
| rs4752028 (C/T; *SHTN1*) | cg00750430 (*SHTN1*) | 1.2 [0.2]; $8.7 \times 10^{-9}$ | 0.13 [0.11]; 0.25 | -16.7; 1 |
| rs4752028 (C/T; *SHTN1*) | cg03968911 (*SHTN1*) | -0.8 [0.1]; $8.7 \times 10^{-9}$ | -0.11 [0.19]; 0.58 | -32.2; 1 |
| rs4752028 (C/T; *SHTN1*) | cg11398452 (*VAX1*) | -0.8 [0.1]; $8.7 \times 10^{-9}$ | -0.11 [0.19]; 0.56 | 30.2; <0.001 |
| rs1258763 (C/T; n/a) | cg04870120 (n/a) | -1.2 [0.3]; $1.3 \times 10^{-6}$ | 0.04 [0.05]; 0.38 | -68.4; 1 |
| rs1873147 (G/A; n/a) | cg04194852 (*TPM1*) | 1.5 [0.3]; $1.5 \times 10^{-8}$ | 0.09 [0.10]; 0.38 | -13.1; 1 |
| rs8076457 (T/C; *NTN1*) | cg18901140 (n/a) | -1.1 [0.2]; $3.0 \times 10^{-7}$ | 0.02 [0.07]; 0.74 | -34.6; 1 |
| rs8076457 (T/C; *NTN1*) | cg19788727 (*NTN1*) | -0.9 [0.2]; $3.0 \times 10^{-7}$ | 0.03 [0.05]; 0.51 | -13.9; 1 |
| rs8076457 (T/C; *NTN1*) | cg02481697 (*NTN1*) | -0.8 [0.2]; $3.0 \times 10^{-7}$ | 0.01 [0.05]; 0.83 | 0.65; 0.01 |
| rs8076457 (T/C; *NTN1*) | cg01862363 (*NTN1*) | -0.6 [0.1]; $3.0 \times 10^{-7}$ | 0.03 [0.09]; 0.78 | 0.11; 0.016 |
| rs8076457 (T/C; *NTN1*) | cg16107528 (*NTN1*) | -0.7 [0.1]; $3.0 \times 10^{-7}$ | -0.01 [0.05]; 0.98 | 4.3; <0.001 |
| rs1808191 (C/A; *PLEKHM1P1*) | cg14501219 (*LOC146880*) | -1.0 [0.2]; $2.9 \times 10^{-6}$ | 0.09 [0.05]; 0.051 | NA* |
| rs1991401 (G/A; *CEP95*)<br>rs1808191 (C/A; | cg02598441 (*LOC146880*) | 0.4 [0.1]; $4.3 \times 10^{-7}$ | -0.04 [0.07]; 0.59 | NA** |
| *PLEKHM1P1*) |  |  |  |  |
| rs3746101<br>(T/G;<br>*MKNK2*) | cg05254098<br>(*MKNK2*) | -1.0 [0.2];<br>$5.0 \times 10^{-6}$ | -0.03 [0.05];<br>0.54 | -37.2; 1 |
| rs3746101<br>(T/G;<br>*MKNK2*) | cg17068236<br>(*MKNK2*) | 0.8 [0.2]; $5.0 \times 10^{-6}$ | -0.02 [0.05];<br>0.58 | -91.0; 1 |

\* This region was too sparsely genotyped to apply the co-localisation analysis
\*\* This CpG had two mQTLs, so we did not apply the co-localisation analysis

### Testing for linkage: joint-likelihood mapping to assess co-localisation

We used a co-localisation method to assess if there was evidence that methylation and liability to nsCL/P are driven by the same causal effect at each locus. Of the 20 CpGs instrumented by a single mQTL, we found evidence for co-localisation at four CpGs (cg11398452, cg01862363, cg02481697 and cg16107528), with three of these being associated with the same mQTL (**Table 1**).

With the addition of the CpG with two mQTLs (cg02598441), we found strongest evidence that methylation at five CpGs are putative mediators of genetic liability to nsCL/P at four SNPs (**Table 1**). Of these four SNPs, three SNPs were available in the imputed GOYA data (rs807647, rs1808191 and rs4752028). Two of the SNPs (intergenic rs8076457 and rs1808191 near *PLEKHM1P1*) consistently replicated as mQTLs in GOYA, with the third SNP rs4752028 replicating as an mQTL for two out of three CpG sites but not the CpG-SNP pairing with evidence of co-localisation (**Supplementary Table 1**).

### Comparison between methylation and gene expression

In a look-up of the four identified SNPs in the GTex and NESDA NTR Conditional eQTL databases, we found strong evidence that rs4752028 at *SHTN1* (which is an mQTL for cg11398452 at *VAX1*) is an eQTL for the nearby *SHTN1* gene (**Table 2**). There was also strong evidence that both rs1808191 at *PLEKHM1P1* and rs1991401 at *CEP95/DDX5* (which are mQTLs for cg02598441 at *LOC146880*) are eQTLs for six nearby genes, including *CEP95* and *DDX5*, which were identified through both databases (**Table 2**). There was no evidence that intergenic SNP rs8076457 (which is an mQTL for cg01862363, cg02481697 and cg16107528 at *NTN1*) is associated with gene expression (**Table 2**).

**Table 2.** Associations with gene expression at identified SNPs in two eQTL databases

| SNP<br>(annotated gene) | CpG<br>(annotated gene) | GTex (gene, tissue, effect size, P-value) | NESDA NTR Conditional eQTL Catalog (gene, tissue, effect size, P-value) |
| --- | --- | --- | --- |
| rs4752028<br>(*SHTN1*) | cg11398452<br>(*VAX1*) | *SHTN1*, whole blood, 0.35, $1.4 \times 10^{-15}$ | *SHTN1*, whole blood, 0.68, $2.7 \times 10^{-159}$ |
| rs8076457<br>(*NTN1*) | cg01862363<br>(*NTN1*) | N/A | N/A |
|  | cg02481697<br>(*NTN1*) |  |  |
|  | cg16107528<br>(*NTN1*) |  |  |
| rs1808191<br>(*CEP95*) | cg02598441<br>(*LOC146880*) | *RP13-104F24.3*, sun exposed skin, -0.31, $1.8 \times 10^{-15}$<br>*SMURF2*, transformed fibroblasts, 0.28,<br>$3.8 \times 10^{-13}$<br>*has-mir-6080*, sun<br>exposed skin, -<br>0.29, $1.5 \times 10^{-11}$<br>*PLEKHM1P*, sun<br>exposed skin, -<br>0.13, $3.2 \times 10^{-5}$ | N/A |
| rs1991401<br>(*PLEKHM1P1*) | | *DDX5*, whole<br>blood, -0.30,<br>$1.8 \times 10^{-19}$<br>*CEP95*, whole<br>blood, 0.14,<br>$2.3 \times 10^{-7}$<br>*MILR1*, whole<br>blood, 0.24,<br>$1.4 \times 10^{-5}$ | *DDX5*, whole<br>blood, -0.22,<br>$6.2 \times 10^{-108}$<br>*CEP95*, whole<br>blood, 0.13,<br>$1.3 \times 10^{-16}$ |

### Comparison to EWAS results

At cg02598441 (*LOC146880*), the direction of effect estimated in our first (forward) MR analysis was concordant with that in the Brazilian EWAS study, with an EWAS P-value (2.4x10^-3^) that survived Bonferroni correction for five tests (**Table 3**). The direction of estimated effect was also concordant between studies at the three CpGs at *NTN1*, but the smallest EWAS P-value was 0.12. At cg11398452 (*VAX1*), the direction of estimated effect was discordant between our MR analysis and the Brazilian EWAS, with a small EWAS P-value (9.4x10^-3^) (**Table 3**).

**Table 3.** Comparison to methylation data in blood samples from children with an orofacial cleft.

| SNP<br>(annotated gene) | CpG<br>(annotated gene) | Forward MR<br>(effect size*<br>[standard error];<br>P-value) | Brazilian<br>nsCL/P<br>EWAS<br>(effect size**<br>[standard error];<br>P-value) | Correlation<br>between<br>blood<br>and lip in<br>the Cleft<br>Collective<br>(correlation<br>coefficient; P-value) | Correlation<br>between<br>blood<br>and palate in<br>the Cleft<br>Collective<br>(correlation<br>coefficient; P-value) | P-value<br>for difference in mean<br>blood DNA<br>methylation<br>between<br>CLO, CLP<br>and CPO<br>in the<br>Cleft<br>Collective |
| --- | --- | --- | --- | --- | --- | --- |
| rs4752028<br>(*SHTN1*) | cg11398452<br>(*VAX1*) | -0.8<br>[0.1];<br>$8.7 \times 10^{-9}$ | 0.01<br>[0.05];<br>$9.4 \times 10^{-3}$ | 0.20;<br>0.057 | 0.07;<br>0.613 | 0.148 |
| rs8076457<br>(*NTN1*) | cg01862363<br>(*NTN1*) | -0.8<br>[0.2];<br>$3.0 \times 10^{-7}$ | -0.030<br>[0.037];<br>$2.1 \times 10^{-1}$ | -0.04;<br>0.699 | -0.02;<br>0.87 | 0.828 |
| | cg02481697<br>(*NTN1*) | -0.6<br>[0.1];<br>$3.0 \times 10^{-7}$ | -0.023<br>[0.024];<br>$1.7 \times 10^{-1}$ | 0.29;<br>0.005 | -0.06;<br>0.684 | 0.286 |
| | cg16107528<br>(*NTN1*) | -0.7<br>[0.1];<br>$3.0 \times 10^{-7}$ | -0.019<br>[0.016];<br>$1.2 \times 10^{-1}$ | 0.34;<br>0.001 | -0.11;<br>0.404 | 0.646 |
| rs1808191<br>(*CEP95*) | cg02598441<br>(*LOC146880*) | 0.4<br>[0.1];<br>$4.3 \times 10^{-7}$ | 0.020<br>[0.007];<br>$2.4 \times 10^{-3}$ | 0.32;<br>0.002 | 0.13;<br>0.359 | 0.548 |
| rs1991401<br>(*PLEKHM1P1*) |  |  |  |  |  |  |

\* Effect size for forward MR can be interpreted as the difference in risk of nsCL/P per standard deviation increase in methylation beta value.
\*\* Effect size for the Brazilian EWAS can be interpreted as the difference in mean methylation beta value in participants with nsCL/P compared to controls.

### Tissue and cleft-subtype-specific variation

At most of the five identified CpGs, methylation in blood, lip and palate tissues was somewhat correlated (correlation coefficients ranging -0.11 to 0.32), particularly between blood and lip tissue. We found weak evidence that mean methylation values in any of the three tissues differed between cleft subtypes. However, the analysis was likely underpowered to detect small to moderate correlations (**Table 3**).

## Discussion

In this study, we employed a framework that aims to identify putative mediators of genetic influences on nsCL/P via DNA methylation. We found five CpG sites, in three independent regions (*VAX1, LOC146880, NTN1*), with either evidence of co-localisation between variants influencing methylation and nsCL/P or evidence of two independent variants affecting both nsCL/P and methylation.

We found lower methylation at the CpG at *VAX1* (cg11398452) in association with the nsCL/P risk allele C of the SNP rs4752028 at *SHTN1*. This SNP is strongly associated with lower expression of *SHTN1* according to two eQTL databases. *VAX1* is a homeobox containing gene that has been shown to be expressed in the developing brain ^44 45^ and SNPs in *VAX1* have been shown to be associated with nsCL/P in multiple independent GWAS across distinct populations ^4 7 8 46 47^. *VAX1* knock-out mice have been shown to develop cleft palate, suggesting *VAX1* has a potentially important role in nsCL/P aetiology ^4^. *SHTN1*, sometimes known as *KIAA1598*, codes for the protein shootin1 that is involved in neuronal polarization^48^ and has also been reported to be relevant to the aetiology of nsCL/P in several studies ^8 49 50^. It is difficult to distinguish the more significant locus between the *VAX1* and *SHTN1* genes because of their close proximity and similar expression profiles in mice and it is unclear which is the functional gene in the area ^8 45 50 51^.

We did not replicate the association between methylation at cg11398452 and rs4752028 in the GOYA data, but we did find that the SNP was strongly associated with methylation at nearby probes. The lack of replication could be because of technical effects, ancestral differences between cohorts or the enrichment of GOYA for overweight and obese mothers, which may introduce selection bias. The direction of association between cg11398452 methylation and nsCL/P was opposite in our MR study compared to a previously published observational EWAS. However, there are several potential explanations for these discordant results, including differences in populations (European vs Brazilian), tissue (cord blood vs whole blood), age of participants (newborns vs children over six years old), or a lack of power giving rise to spurious associations in either analysis.

We found higher methylation at the CpG at *LOC146880* (cg02598441) in association with the G allele of rs1991401 in *DDX5* and the C allele of rs1808191 in *PLEKHM1P*. rs1991401 in *DDX5* was associated with reduced expression of *DDX5* and increased expression of *CEP95* and *MILR1* while rs1808191 in *PLEKHM1P* was associated with increased expression of *SMURF2* but decreased expression of *PLEKHM1P*, *RP13-104F24.3* and *has-mir-6080*. However, there was weak evidence that the SNPs affected expression of the same genes. *DDX5* is involved in RNA helicase processes that are highly relevant to important cellular processes while *PLEKHM1P* and *LOC146880* are pseudogenes ^45^. There is no robust evidence from previous literature to support an association between genetic variation of these genes and nsCL/P. The SNP in *PLEKHM1P* replicated as an mQTL in the GOYA dataset but there was not sufficient data to test the SNP in *DDX5*.

We found lower methylation at three CpGs at *NTN1* in association with the nsCL/P risk allele T of the SNP rs8076457, an intergenic SNP close to *NTN1*. rs8076457, the mQTL for cg08162363, cg02481697 and cg16107528, did not robustly associate with gene expression levels in two datasets. The function of *NTN1* is still largely unknown but is thought to be involved in cell migration during development ^45^. *NTN1* has been previously discussed as a strong candidate gene for nsCL/P ^12^; *NTN1* may affect liability to nsCL/P via epistatic interactions, there is some evidence that *NTN1* knock-out mice show consistency with the cleft palate phenotype and *NTN1* expression is localised to the palate ^12 52^. rs8076457 replicated as an mQTL across all relevant CpGs in the GOYA dataset.

Previous work has identified many functional possibilities for genetic risk variants for nsCL/P ^14 52 53^ but this study is the first to specifically look at the role of DNA methylation in mediating genetic susceptibility to nsCL/P. Additional strengths of this study include the integration of multiple data sources, for example ALSPAC, which provided access to detailed phenotype, genotype and epigenetic data. The nsCL/P GWAS summary statistics allowed a comprehensive genome-wide analysis in a large dataset. The Brazilian cohort EWAS results allowed a comparison of the influence of methylation on nsCL/P according to observational and MR studies. The use of the GOYA replication cohort, allowed triangulation of evidence for mQTLs across different studies. Finally, the Cleft Collective data allowed us to compare genome-wide DNA methylation in different tissues and subtypes of cleft.

There are, several limitations to this study. First, methylation and expression in the studied tissues (postnatal cord blood, whole blood, lip and palate tissue) may not accurately reflect that in the developing orofacial tissue where epigenetic processes could feasibly influence susceptibility to nsCL/P. However, a previous study has identified a high correlation between blood and lip tissue, both taken at the time of first surgery in a UK cohort of patients with non-familial nsCL/P ^15^. Previous analysis looking for tissue-specific signals for nsCL/P did not find evidence of enrichment and concluded that this may be due to tissue type differences ^2^. Second, cleft lip only (CLO) and cleft lip and palate (CLP) cases were analysed together as one group in the GWAS, MR analyses and the previously published EWAS. Increasingly, evidence suggests that these subtypes are molecularly and aetiologically distinct and should be analysed separately ^2 3^, but we were limited by the data available from previous studies. Although we found no evidence of differential methylation between subtypes at our five identified CpGs, there may be other loci where methylation mediates genetic influences on more specific cleft subtypes. Third, although efforts were made to select only non-syndromic cases for the Cleft Collective analysis, we cannot guarantee that no syndromic cases were included, and children with syndromes may have very different methylation profiles. Fourth, a major limitation of this study is that some of the steps, particularly the reverse MR, are likely to be statistically underpowered. Fifth, as the majority of mQTLs were instrumented by just a single genetic variant, we were unable to distinguish between mediation and horizontal pleiotropy and therefore proposed mediation is putative. Finally, although mQTLs were largely concordant between ALSPAC and GOYA, the mQTL (in *SHTN1*) found to co-localise with liability to nsCL/P in ALSPAC did not replicate in GOYA.

In conclusion, we identified three putative loci where DNA methylation may mediate genetic susceptibility to nsCL/P. Future work, determining the function of these genes and the epigenetic modulation of their expression relevant to prenatal orofacial development could provide important aetiological insights.

## Acknowledgements

ALSPAC: We are extremely grateful to all the families who took part in this study, the midwives for their help in recruiting them, and the whole ALSPAC team, which includes interviewers, computer and laboratory technicians, clerical workers, research scientists, volunteers, managers, receptionists, and nurses. We would like to acknowledge Tom Gaunt, Oliver Lyttleton, Sue Ring, Nabila Kazmi, and Geoff Woodward for their earlier contribution to the generation of ARIES data (ALSPAC methylation data).

Cleft Collective: This publication (project number CC002) involves data derived from independent research funded by The Scar Free Foundation (REC approval 13/SW/0064). We are grateful to the families who participated in the study, the UK NHS cleft teams, and the Cleft Collective team, who helped facilitate the study. We also thank the Bristol Bioresource Laboratory team for the sample processing and storage. Special thanks go to Kerry Humphries, Amy Davies and Karen Ho.

GOYA: GOYA is a study nested within the Danish National Birth Cohort. The authors want to thank the many pregnant women who have taken part in the cohort and the midwives who collected the umbilical cord samples. Additionally, we would like to acknowledge Thorkild I.A. Sørensen for his contribution to the generation of the GOYA data.

## Supplementary Material

### nsCL/P GWAS methods

The transmission disequilibrium test (TDT) ^1^ evaluates the frequency with which parental alleles are transmitted to affected offspring and is a family based association test of genetic linkage in the presence of genetic association. The TDT was run on 638 parent-offspring trios and 178 parent-offspring duos of European, descent, publicly available from dbGAP, to determine genome-wide genetic variation associated with nsCL/P. GWAS genotypes and phenotypes available at dbGaP (https://www.ncbi.nih/gov/gap; accession number phs000094.v1.p1).

The Bonn-II study ^2^ summary statistics from a case-control GWAS of 399 nsCL/P cases and 1,318 controls were meta-analysed using a fixed effect inverse-variance weighted method, in terms of effect size and standard error, with the TDT GWAS summary statistics using METAL ^3^ based on a previously described protocol for combining TDT and case-control studies ^4^. The final sample including 1215 cases and 2772 controls.

### Avon Longitudinal Study of Parents and Children (ALSPAC)

To identify mQTLs (SNPs associated with DNA methylation), we used data from the Avon Longitudinal Study of Parents and Children (ALSPAC). ALSPAC is a longitudinal study that recruited pregnant women living in the former county of Avon (UK) with expected delivery dates between 1 April 1991 and 31 December 1992^5 6^. Written, informed consent was obtained for all participants. Ethics approval for the study was obtained from the ALSPAC Ethics and Law Committee and the Local Research Ethics Committee. The study website contains details of all available data through a searchable data dictionary (http://www.bristol.ac.uk/alspac/researchers/dataaccess/datadictionary/). In addition to collecting detailed questionnaire and clinic data for the whole cohort, the study has generated genome-wide DNA methylation and genotype data for subsets.

As part of the Accessible Resource for Integrated Epigenomic Studies (ARIES) project ^7^, genome-wide DNA methylation data were generated for 1018 ALSPAC mother-child pairs across five time-points. For the purposes of the current study, we used data generated using offspring cord blood samples collected at birth. Generation of these data are described in detail elsewhere. Briefly, DNA samples were bisulfite treated and DNA methylation was quantified using the Illumina Infinium HumanMethylation450K BeadChip assay, which measures DNA methylation at over 480,000 CpG sites across the genome. After quality control and functional normalisation using the r package meffil ^8^, data were reported as methylation beta values, ranging from 0 (completely unmethylated) to 1 (completely methylated). Genotype data were available for all 1018 children in ARIES, generated using the Illumina HumanHap550 quad genome-wide SNP genotyping platform. Individuals were excluded from further analysis based on having incorrect gender assignments, minimal or excessive heterozygosity (0.345 for the Sanger data and 0.330 for the LabCorp data), disproportionate levels of individual missingness (>3%), evidence of cryptic relatedness (>10% IBD) and being of non-European ancestry (as detected by a multidimensional scaling analysis seeded with HapMap 2 individuals.

ALSPAC methylation and genotype data have previously been used to generate a database of mQTLs (http://www.mqtldb.org/) ^9^. The database contains summary statistics for all mQTLs with a P-value <1*10-5 for the association between SNP and CpG. For part of our study, we required specific CpG-SNP associations that were unavailable from mQTLdb.org. Therefore, for required CpGs, we replicated the methods in the original study: we excluded individuals with missing genotype or covariate data, leaving 787 children. We then rank-normalised the methylation data to remove outliers and then controlled for covariates, potential batch effects and the influence of cell heterogeneity by regressing data points on sex, the first 10 ancestry principal components, bisulfite-converted DNA batch and blood cell proportions ^10 11^estimated using the Houseman method. The residuals were then used as the outcome variable in a linear regression model in PLINK^12^ to calculate the relevant CpG-SNP associations.

### Genetics of Overweight Young Adults (GOYA)

The Genetics of Overweight Young Adults (GOYA) study is described previously by Paternoster et al ^13^. It is based on the Danish National Birth Cohort that included 92,000 pregnant women and their pregnancies during 1996-2002. Of 67,853 women who had given birth to a live born infant, had provided a blood sample during pregnancy and had BMI information available, 3.6% of these women with the largest residuals from the regression of BMI on age and parity (all entered as continuous variables) were selected for GOYA. The BMI for these 2451 women ranged from 32.6 to 64.4. From the remaining cohort a random sample of similar size (2450) was also selected. DNA methylation data were generated for the offspring of 1000 mothers in the GOYA study. I.e. “cases” had mothers with a BMI>32 and “controls” were sampled from the normal BMI distribution (can include mothers with a BMI>32).

Methylation data were generated at the University of Bristol as described above for ALSPAC. Data were QCd and normalised using the meffil R package ^8^. Genome-wide genotyping on the Illumina 610k quad chip was carried out at the Centre National de Genotypage, Evry, France. Individuals were excluded from further analysis based on having incorrect gender assignments, minimal or excessive heterozygosity (>35% or <30.2%), disproportionate levels of individual missingness (>5%), relatedness and being of non-European ancestry (as detected by a multidimensional scaling analysis seeded with HapMap 2 individuals.

### Replication of mQTL results

**Supplementary Table 1:**
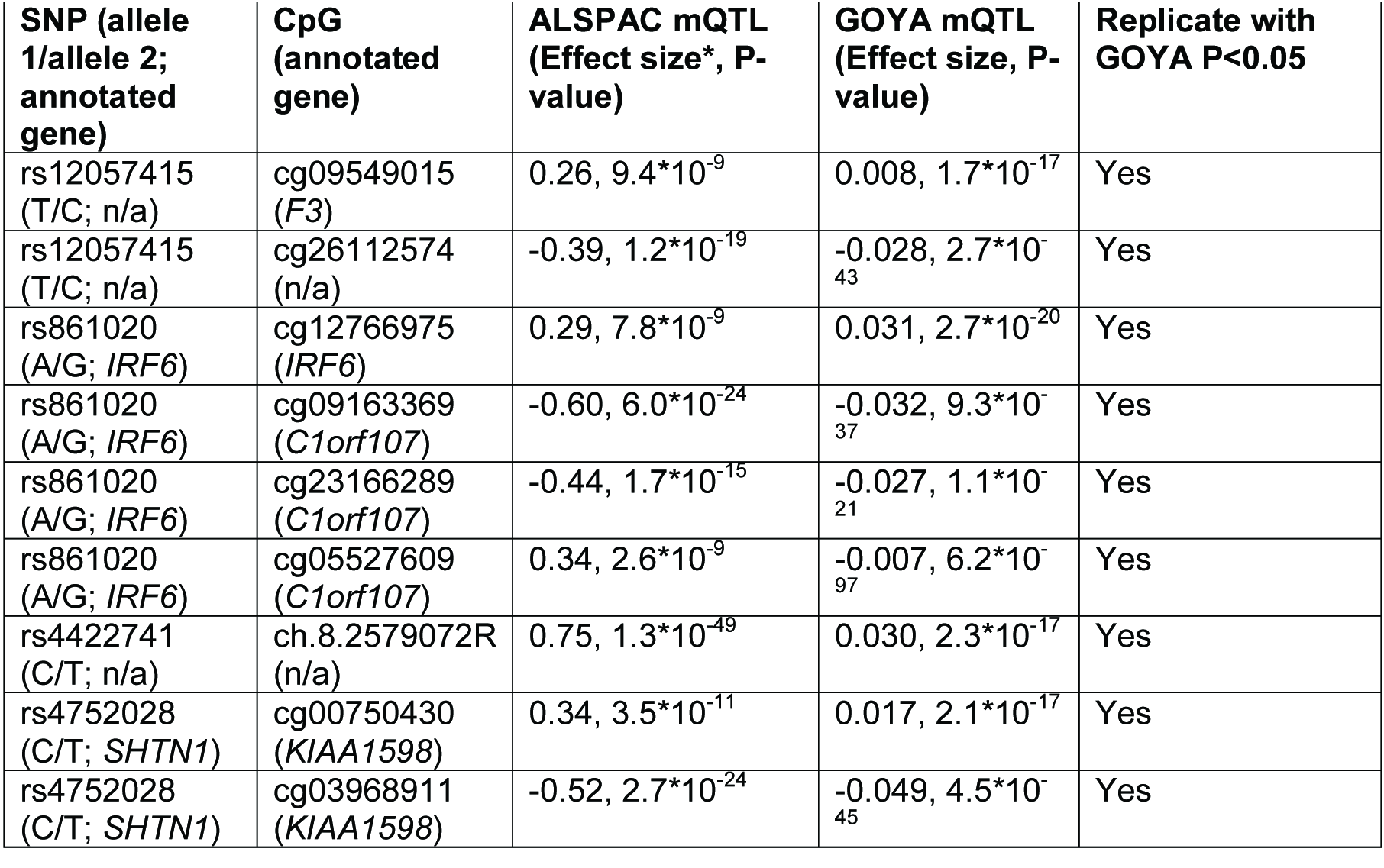
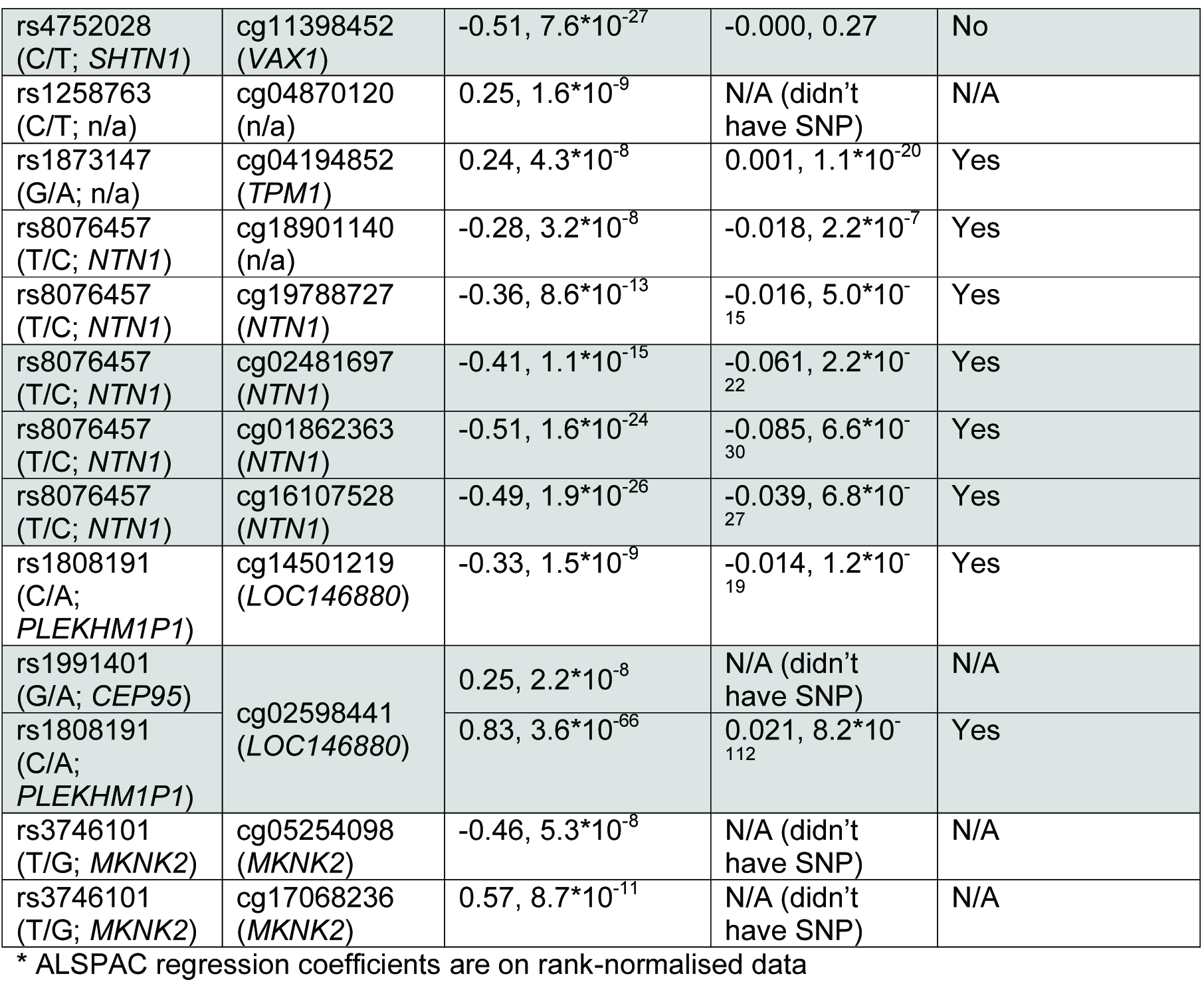
mQTL replication

| SNP (allele 1/allele 2; annotated gene) | CpG (annotated gene) | ALSPAC mQTL (Effect size*, P-value) | GOYA mQTL (Effect size, P-value) | Replicate with GOYA P<0.05 |
| --- | --- | --- | --- | --- |
| rs12057415 (T/C; n/a) | cg09549015 (F3) | 0.26, $9.4 \times 10^{-9}$ | 0.008, $1.7 \times 10^{-17}$ | Yes |
| rs12057415 (T/C; n/a) | cg26112574 (n/a) | -0.39, $1.2 \times 10^{-19}$ | -0.028, $2.7 \times 10^{-43}$ | Yes |
| rs861020 (A/G; *IRF6*) | cg12766975 (*IRF6*) | 0.29, $7.8 \times 10^{-9}$ | 0.031, $2.7 \times 10^{-20}$ | Yes |
| rs861020 (A/G; *IRF6*) | cg09163369 (*C1orf107*) | -0.60, $6.0 \times 10^{-24}$ | -0.032, $9.3 \times 10^{-37}$ | Yes |
| rs861020 (A/G; *IRF6*) | cg23166289 (*C1orf107*) | -0.44, $1.7 \times 10^{-15}$ | -0.027, $1.1 \times 10^{-21}$ | Yes |
| rs861020 (A/G; *IRF6*) | cg05527609 (*C1orf107*) | 0.34, $2.6 \times 10^{-9}$ | -0.007, $6.2 \times 10^{-97}$ | Yes |
| rs4422741 (C/T; n/a) | ch.8.2579072R (n/a) | 0.75, $1.3 \times 10^{-49}$ | 0.030, $2.3 \times 10^{-17}$ | Yes |
| rs4752028 (C/T; *SHTN1*) | cg00750430 (*KIAA1598*) | 0.34, $3.5 \times 10^{-11}$ | 0.017, $2.1 \times 10^{-17}$ | Yes |
| rs4752028 (C/T; *SHTN1*) | cg03968911 (*KIAA1598*) | -0.52, $2.7 \times 10^{-24}$ | -0.049, $4.5 \times 10^{-45}$ | Yes |
| rs4752028<br>(C/T; *SHTN1*) | cg11398452<br>(*VAX1*) | -0.51, $7.6 \times 10^{-27}$ | -0.000, 0.27 | No |
| rs1258763<br>(C/T; n/a) | cg04870120<br>(n/a) | 0.25, $1.6 \times 10^{-9}$ | N/A (didn't have SNP) | N/A |
| rs1873147<br>(G/A; n/a) | cg04194852<br>(*TPM1*) | 0.24, $4.3 \times 10^{-8}$ | 0.001, $1.1 \times 10^{-20}$ | Yes |
| rs8076457<br>(T/C; *NTN1*) | cg18901140<br>(n/a) | -0.28, $3.2 \times 10^{-8}$ | -0.018, $2.2 \times 10^{-7}$ | Yes |
| rs8076457<br>(T/C; *NTN1*) | cg19788727<br>(*NTN1*) | -0.36, $8.6 \times 10^{-13}$ | -0.016, $5.0 \times 10^{-15}$ | Yes |
| rs8076457<br>(T/C; *NTN1*) | cg02481697<br>(*NTN1*) | -0.41, $1.1 \times 10^{-15}$ | -0.061, $2.2 \times 10^{-22}$ | Yes |
| rs8076457<br>(T/C; *NTN1*) | cg01862363<br>(*NTN1*) | -0.51, $1.6 \times 10^{-24}$ | -0.085, $6.6 \times 10^{-30}$ | Yes |
| rs8076457<br>(T/C; *NTN1*) | cg16107528<br>(*NTN1*) | -0.49, $1.9 \times 10^{-26}$ | -0.039, $6.8 \times 10^{-27}$ | Yes |
| rs1808191<br>(C/A; *PLEKHM1P1*) | cg14501219<br>(*LOC146880*) | -0.33, $1.5 \times 10^{-9}$ | -0.014, $1.2 \times 10^{-19}$ | Yes |
| rs1991401<br>(G/A; *CEP95*) | cg02598441<br>(*LOC146880*) | 0.25, $2.2 \times 10^{-8}$ | N/A (didn't have SNP) | N/A |
| rs1808191<br>(C/A; *PLEKHM1P1*) | | 0.83, $3.6 \times 10^{-66}$ | 0.021, $8.2 \times 10^{-112}$ | Yes |
| rs3746101<br>(T/G; *MKNK2*) | cg05254098<br>(*MKNK2*) | -0.46, $5.3 \times 10^{-8}$ | N/A (didn't have SNP) | N/A |
| rs3746101<br>(T/G; *MKNK2*) | cg17068236<br>(*MKNK2*) | 0.57, $8.7 \times 10^{-11}$ | N/A (didn't have SNP) | N/A |

\* ALSPAC regression coefficients are on rank-normalised data

